# Timed sulfonylurea modulation improves locomotor and sensory dysfunction following spinal cord injury

**DOI:** 10.1101/2024.05.06.592741

**Authors:** Guo-Ying Xu, Manjit Maskey, Zizhen Wu, Qing Yang

## Abstract

Traumatic injury to the spinal cord (SCI) results in immediate necrosis and delayed secondary expansion of neurological damage, often resulting in lifelong paralysis, neurosensory dysfunction, and chronic pain. Progress hemorrhagic necrosis (PHN) and excessive excitation are the primary sources of neural injury triggered by various insults, causing neuronal cytotoxicity and the gradual enlargement of lesions. Recent approaches have involved blocking TRPM4, a contributor to PHN, using the sulfonylurea (SUR) subunits regulator glibenclamide. However, since SUR subunits are expressed in both neurons and glial cells in the spinal cord and sensory neurons, forming functional K_ATP_ channels, the use of glibenclamide can exacerbate the development of SCI-induced chronic pain. In this study, we explored a treatment strategy involving the administration of glibenclamide, which suppresses PHN, and diazoxide, which protects against neuronal excitation and inflammation, at different time intervals post-SCI. Our goal is to determine whether this approach significantly enhances both sensory and motor function. Contusive SCI was induced at spinal segment T10 in adult male and female rats. Immunostaining, electrophysiological recording *in vitro*, spectrophotometric assay, Western blot, and behavioral tests were performed to detect the outcomes. Results showed that the timed application of glibenclamide and diazoxide at various post-SCI time points significantly improved locomotor function and mitigated the development of SCI-induced chronic pain. These preclinical studies introduce a promising treatment strategy for addressing SCI-induced dysfunction.

## Introduction

The spinal cord, located within the vertebrae, plays a vital role in transmitting sensory, motor, and autonomic signals between the brain and the rest of the body. Unfortunately, it can sustain injuries from sudden traumatic impacts to the spine, often resulting from accidents or violence. This leads to an annual occurrence of 250,000 to 500,000 new cases of spinal cord injury (SCI) worldwide, with victims typically experiencing permanent loss of locomotor and sensory function below the injury site, along with the development of chronic pain ^1^. To date, there is no effective treatment available for SCI. The primary injury to the spinal cord usually causes partial damage to blood vessels and neuronal tissues, leaving many neuronal axons and interneurons temporarily intact ^1^. However, these surviving tissues often undergo progressive neuronal necrosis and lesion expansion due to secondary degeneration. So far, various signaling pathways involved in the secondary cascades are identified ^2^, which has opened possibilities for intervening in the early stages of SCI to reduce lesion expansion and enhance both sensory and motor functions.

Progressive hemorrhagic necrosis (PHN) is a common occurrence in the secondary injury following SCI ^3^. Abundant expression of TRPM4, which forms complexes with regulatory sulfonylurea 1 (SUR1), has been shown to initiate PHN in capillary endothelial cells after SCI ^4, 5^. Notably, secondary hemorrhage and lesion expansion are significantly reduced in both Sur1 mutant mice ^6^ and Trpm4 null mice ^5^ after SCI. Additionally, pharmacological blockade of TRPM4 channels has been found to decrease PHN and enhance motor function recovery ^3, 4^. However, it’s worth noting that blocking TRPM4 through its regulatory sulfonylurea (SUR) subunits could have potential drawbacks. SUR subunits are not only upregulated in capillary endothelial cells following SCI but are also expressed in neurons, glial cells, and the mitochondrial membrane in the spinal cord to form functional K_ATP_ channels (an octomeric complex consisting of Kir6.1 or Kir6.2 subunits and SUR). For instance, SUR1 associates with Kir6.2 to form functional K_ATP_ channels in neurons ^7, 8^. In contrast, Kir6.1 and SUR2 are primarily distributed in astrocytes within the spinal cord ^9^. Consequently, inhibiting SUR subunits may potentially lead to neuronal hyperexcitability and glial activation. While blocking SUR1 with glibenclamide immediately after spinal contusion for 24 hours decreases hemorrhagic necrosis and improves long-term locomotion and coordination ^3, 10, 11^, it worsens sensory function after SCI, likely by influencing glial cell activation ^12^. Conversely, diazoxide, a specific K_ATP_ channel opener, mitigates damage-induced neurodegeneration and glial activation through K_ATP_ channel-mediated mitochondrial and cellular protection ^13–15^. Therefore, an approach that can reduce PHN while preserving the beneficial effects of opening K_ATP_ channels may yield better outcomes for both motor and sensory function compared to glibenclamide or diazoxide used alone.

In this investigation, we manipulated the activity of SUR subunits using their agonists or inhibitors to regulate TRPM4 or K_ATP_ channel activity at distinct time points post-SCI, aiming to evaluate the effect of the treatment strategy. The findings revealed that the timed administration of glibenclamide and diazoxide at different periods after SCI significantly enhanced locomotor function and alleviated the progression of chronic pain induced by SCI. These preclinical studies present a promising therapeutic approach for managing dysfunction resulting from spinal cord injury.

## Methods

Our procedures adhered to the guidelines set forth by the National Institutes of Health and received approval from the Institutional Animal Care and Use Committee at the University of Texas Medical Branch in Galveston. We acquired adult Sprague-Dawley rats weighing between 200-250g from Charles River Laboratories (Wilmington, MA). Both male and female rats were included in the study, and no noticeable differences related to sex were observed.

The animals were housed two per cage in a temperature-controlled room with a 12-hour light/dark cycle, maintaining a temperature of 21 ± 1 °C. They had unrestricted access to food and water throughout the study. Prior to the commencement of the experiments, the rats were allowed to acclimate to their surroundings for one week.

### Spinal cord injury (SCI)

SCI in humans typically arises from spinal cord contusion. To replicate this condition, we employed a spinal cord contusion model. Recognizing the variability in the extent of spinal cord damage resulting from different contusion models ^16^, we chose to induce the spinal cord contusion by the Infinite Horizon impactor (Precision Systems & Instrumentation, Lexington, KY) since this device reliably simulates a contusive injury by rapidly applying a precisely defined force ^17^.

The experimental animals were anesthetized using a combination of acepromazine (0.75 mg/kg), xylazine (20 mg/kg), and ketamine (80 mg/kg). A laminectomy was performed, followed by the induction of a moderate spinal contusion at the T10 level using a force of 150 kdyne and a 1-second dwell time. In contrast, the sham group only had the backbone removal without the spinal cord injury. Afterward, the muscles were stitched over the backbone, and the skin was closed with clips. The animals that underwent SCI/laminectomy were then placed in cages equipped with heating pads set at around 37°C. Animals received analgesic injections (buprenorphine; 0.02 mg/kg, i.p.) twice daily for a duration of 5 days, along with prophylactic antibiotics (Baytril, 2.5 mg/kg, i.p.) administered for 10 days. Manual bladder evacuation was performed twice daily until signs of bladder function recovery were observed. Regular monitoring of the rats was carried out to detect any signs of severe pain, such as excessive grooming, pronounced inactivity, or self-mutilation. Any animals that showed severe signs of pain were euthanized right away.

### DRG neuron dissociation and culturing

The SCI rats underwent deep anesthesia with Beuthanasia (75 mg/kg, Merck Animal Health, Kenilworth, NJ), followed by intracardial perfusion of cold phosphate-buffered saline (PBS). The ganglia were then dissected from lumbar vertebrae 4 and 5. The L4/L5 DRGs were minced with a fine scissor and placed in Dulbecco’s Modified Eagle Medium (DMEM; Invitrogen, Waltham, MA) containing 0.35 mg/mL trypsin (Worthington, Lakewood, NJ) and 0.8 mg/mL collagenase (Roche, Mannheim, Germany) for 40 minutes at 34°C. Following this, the cell-containing solution underwent centrifugation (300 rpm for 5 minutes) and resuspension. The isolated cells were then plated onto 8 mm cover glasses (Warner Instruments, Hamden, CT) precoated with 50 mg/mL poly-L-lysine and incubated (37°C, 5% CO2) in DMEM without serum for overnight.

### Electrophysiological recordings

Electrodes were fashioned from borosilicate glass capillaries (BF150-110-10, Sutter Instrument Co., Novato, CA) using a P-1000 micropipette puller (Sutter Instrument Co., Novato, CA), ensuring a resistance of approximately 2 MΩ for subsequent experiments. Neuronal observation was facilitated using an Eclipse Ti inverted microscope (Nikon, Tokyo, Japan). Whole-cell current clamp was conducted utilizing a MultiClamp 700B amplifier with an Axon Digidata 1550B data acquisition system (Molecular Devices, San Jose, CA). The resting membrane potential (RMP) and any spontaneous firing of DRG neurons were recorded by patching the cells at 0 pA. Signals were sampled at a rate of ≥ 10 kHz. The pipette solution comprised (in mM): 134 KCl, 1.8 CaCl_2_, 13.2 NaCl, 1.6 MgCl_2_, 9 HEPES, 3 EGTA, 1 Mg-ATP, and 0.3 Na-GTP (pH 7.2, 300 mOsM). The bath solution included (in mM): 140 NaCl, 1.8 CaCl_2_, 3 KCl, 2 MgCl_2_, 10 glucose, and 10 HEPES (pH 7.4, 320 mOsM).

### Delivery of glibenclamide and/or diazoxide

It has been reported that blocking SUR1 with glibenclamide immediately after a spinal contusion for a 24-hour duration reduces hemorrhagic necrosis and leads to long-term enhancements in locomotion and coordination ^3, 10, 11^. Therefore, glibenclamide is administered acutely (5 mg) beginning 30 minutes after the impact using an intradermally implanted mini-osmotic pump (Alzet Corp., Cupertino, CA) for a 24-hour period. This timing aligns with the fact that hemorrhage is most pronounced between 3 to 24 hours after the acute SCI ^18^, Subsequently, animals received systemic diazoxide treatment for ten consecutive days by intraperitoneal injection, commencing three days after the contusion.

### Behavioral tests

*Reflex sensitivity tests:* The behavioral tests were assessed during the light phase both before and after SCI. The personnel conducting the behavioral tests were unaware of the drug treatments administered. To acclimatize each animal to the testing chamber, they spent 20 minutes in it before the experiments commenced. We employed a series of calibrated *von Frey* filaments (Stoelting, Wood Dale, IL), which were applied to the hairless surface of the hind paws. Mechanical sensitivity thresholds were determined using the “up-down” method ^19^.

*Conditioned place preference (CPP) test for* spontaneous *pain:* For the evaluation of spontaneous pain, we employed a conditioned place preference (CPP) device (Med Associates Inc.,VT) ^20, 21^. This CPP device consisted of one white and one black chambers. During the initial day, each animal underwent a 30-minute habituation period in the CPP chambers. Over the subsequent three days, animals were conditioned with twice-daily injections of analgesic and vehicle, each associated with a specific chamber. In the analgesic conditioning session, the rat received an injection of retigabine (10 mg/kg, i.p.) and was placed in the white chamber for one hour, beginning five minutes after the injection. During a distinct conditioning session, the identical rat was isolated within the black chamber for a duration of one hour subsequent to receiving a saline injection (i.p.). On the fifth day, the test subject, without receiving any injections, was positioned inside the black chamber and allowed unrestricted access to both chambers. The system monitored the duration of time the subject spent in black and white chambers over a 30-minute recording.

*Basso, Beattie, and Bresnahan (BBB) locomotor scoring:* To assess the impact of SCI on locomotor function, we employed the 21-point Basso, Beattie, Bresnahan (BBB) locomotor test ^22^. Subjects were positioned within an open field container, and their locomotor behavior was meticulously observed and evaluated under white light conditions. The testing procedure was carried out daily for the initial five days and subsequently on a weekly basis for a period ranging from 5 to 6 weeks post-contusion. Rats that exhibited a BBB score of 1 or higher on the first day following contusion were excluded for analysis.

*Horizontal ladder:* One week prior to the injury, a training regimen was implemented. During this training, rats were daily coached to traverse a 1-meter-long horizontal ladder with a testing section spanning 0.8 meters.

Within the testing area, there were 10 rungs randomly spaced between 3 to 8 centimeters apart. Each training session encompassed three ladder crossings. On the conclusive day of the training period, the tests were recorded on video. These videotapes were later reviewed in slow motion to enable the quantification of hindlimb misses and footslips throughout the crossings. This evaluation allowed for the assignment of a score to each tested animal. A score of 1 was allocated when an entire paw descended below a rung during testing.

Subsequently, this assessment was repeated once a week for a total of 5 weeks, starting on day 14 following the contusion injury.

### Spectrophotometric Assay for spinal cord hemorrhage

The hemoglobin content in the spinal cord following SCI was quantified using a spectrophotometric assay, following a previously described method ^23^. Rats were euthanized using Beuthanasia (Merck, Kenilworth, NJ) and subsequently subjected to perfusion with cold heparinized Phosphate-buffered saline (PBS) to remove intravascular blood. Spinal cord segments spanning the lesion site (5 mm) were dissected. To each sample, 250 μl of distilled water was added, followed by 30 seconds of homogenization (Fisher Scientific) and 1 minute of sonication (Qsonica, Newtown, CT) on ice. The homogenized samples were then centrifuged at 13,000 rpm for 30 minutes, and the supernatant containing hemoglobin was collected.

For the quantification, 80 µl of Drabkin’s reagent (RICCA Chemical, 266016) was added to 20 µl sample and incubated for 15 minutes to fully convert hemoglobin to cyanomethemoglobin. This product was then assessed using iMark Microplate Reader (Bio-Rad) at ≈540 nm wavelength (OD1). As an additional control measure, blood was obtained from a naïve rat via cardiac puncture after anesthesia. 1 µl blood was then added to 20 µl spinal cord specimen collected from the naïve rat for measurement (OD2). The optical density of all samples from the experimental groups was normalized to that from blooded naïve specimen. The blood volume (µl) from each experimental spinal cord following SCI was calculated as: (OD1/OD2) x (250/20).

### Fluorescence Immunostaining

The naïve rats underwent deep anesthesia with Beuthanasia, followed by perfusion with 4% paraformaldehyde (PFA). Afterward, the L4 and L5 segments of the DRGs from each animal were isolated and placed into 1.5 ml Eppendorf tubes. The DRGs were further fixed with 4% paraformaldehyde in PBS for 3 hours, followed by cryoprotection in 30% sucrose overnight at 4°C. DRGs were then sectioned into 14 µm slices using a cryostat (Epredia, Kalamazoo, MI). Sectioned tissues were then incubated with 5% NGS and 0.5% Triton X-100 followed by primary antibodies targeting Nav1.8 (ASC016, Alomone Labs) and Kir6.2 (ab307371, Abcam). After secondary antibodies incubation and PBS rinsing, the samples were covered with Vectashield and imaged using a Nikon A1 Confocal Laser Microscope System.

### Western Blotting

After the completion of final behavioral tests, the animals underwent deep anesthesia with Beuthanasia followed by perfusion with ice-cold PBS. Subsequently, the L4 and L5 segments of the spinal cords from each rat were extracted and placed into 1.5 ml Eppendorf tubes on dry ice. The tissues were homogenized in 500 μl of RIPA lysis buffer (Sigma, Cat# R0278), to which a protease inhibitor cocktail (Fisher Scientific, Cat# PI78430) was added. Following homogenization, the samples underwent three rounds of sonication (10-second pulses) and were then centrifuged (14,000 rpm for 10 minutes) at 4°C. The total protein concentration of the lysates was measure by Nanodrop One (thermo scientific, Waltham, MA). The samples were then prepared for SDS-PAGE (Bio-Rad, 4-20% Tris-HCl) by diluting them in a 1:1 ratio with Laemmli sample buffer, and 30 μg of protein was loaded into each well. Following electrophoresis, the gel was transferred onto a PVDF membrane using Turbo Transfer system (Bio-Rad, Hercules, CA) and blocked with 5% nonfat dry milk in TBS + 0.1% Tween 20. Membrane was then incubated overnight at 4°C with antibodies against GFAP (Millipore, AB5541), Iba-1 (Wako, 019-19741), and β-actin (Abcam, MA; ab8226). The membrane was then incubated with HRP-conjugated anti-rabbit or anti-mouse IgG (Jackson ImmnuoResearch, PA) for 1 hour at room temperature and developed using the ECL kit (Pierce). Protein expression levels were quantified by measuring optical density using Image J software (NIH). Molecular weight standards were run on each gel, and β-actin served as the loading control.

### Eriochrome cyanine (EC) histochemistry

To distinguish between gray and white matter in spinal cord sections, an Iron-eriochrome cyanine R (EC) staining procedure was employed ^24, 25^. After the completion of all behavioral tests, which occurred 35 or 42 days post-contusion, the animals were humanely euthanized using Beuthanasia (Merck, Kenilworth, NJ) and subsequently subjected to perfusion with cold Phosphate-buffered saline (PBS), followed by a 4% paraformaldehyde solution (PFA). The spinal cords, with the lesion site positioned centrally (10 mm in length), were then extracted and post-fixed in 4% PFA overnight. Afterward, the tissues were immersed in 30% sucrose at 4 °C. The spinal cords at the injury site were encased in cryo-embedding medium (OCT, Sakura Finetek, Torrance, CA) and preserved at a temperature of -80°C. Following this, the tissues were sliced into transverse sections by HM525 NX Cryostat (Epredia, Kalamazoo, MI), each with a thickness of 20 μm. A separation of 200 μm was maintained between these sections. The particular section for each animal that exhibited the most substantial cavity was designated as the lesion epicenter. The assessment of the lesion volume encompassed a span extending from 5 mm caudally to rostrally starting from the epicenter. These sections were subjected to a sequence of procedures, including dehydration, clearing, rehydration, and staining. The stained sections were scrutinized utilizing a Nikon microscope and subjected to analysis with the NIS Elements software.

### Statistics

An individual not directly engaged in the experiment’s execution prepared reagents and labeled them with numerical identifiers. The animals were randomly allocated to different groups. All evaluations of behavior and histological examinations were carried out in a manner that maintained blinding. The un-blinding procedure commenced solely after data collection, data input, and database formatting were concluded. For data analysis, we utilized both Sigmaplot (version 14, Systat software, San Jose, CA) and Prism (version 9.0, GraphPad, La Jolla, CA). The data was presented either as the mean ± standard error of the mean (S.E.M.) or as percentages. A significance level (α) of 0.05 was established for all statistical tests. When comparing more than two groups, we applied either one-way or two-way ANOVA, followed by Tukey’s multiple comparison tests. For comparisons within the same cohort before and after treatment, two-tailed paired t-tests were conducted.

Statistical significance was ascertained for findings with p-values below 0.05, indicated by * for p < 0.05, ** for p < 0.01, and *** for p < 0.001. Comprehensive statistical details for all experiments can be found within the figure captions.

## Results

### 1. Diazoxide decreases the excitability of primary sensory neurons

After spinal cord injury (SCI), the heightened activity of primary sensory neurons plays a crucial role in chronic pain development and maintenance ^21, 26, 27^, we thus first detected the expression of K_ATP_ channels in DRG neurons and its ability for controlling cell excitability. Consistent to what observed by other labs ^7, 8^, Kir6.2 is widely expressed in DRG neurons, including small sized cells expressing Nav1.8 (**Fig. 1A**). Since potassium (K^+^) channels are pivotal in regulating neuronal excitability, we initially investigated whether diazoxide could mitigate the excitability of dorsal root ganglia (DRG, L4/L5) neurons dissociation from SCI rats. Dissociated DRG neurons were subjected to whole-cell current clamp recording (0 pA). Initially, DRG neurons were recorded for 15 seconds in a normal external solution, followed by the local application of diazoxide (100 μM) for 1 minute. The majority of cells exhibited a more negative membrane potential after diazoxide compared to that after vehicle (**Fig. 1B and C**), and the firing rate of the neurons was greatly decreased by diazoxide (**Fig. 1B**). These results indicate that diazoxide can effectively decreases the excitability of DRG neurons induced by SCI.

**Figure 1.**
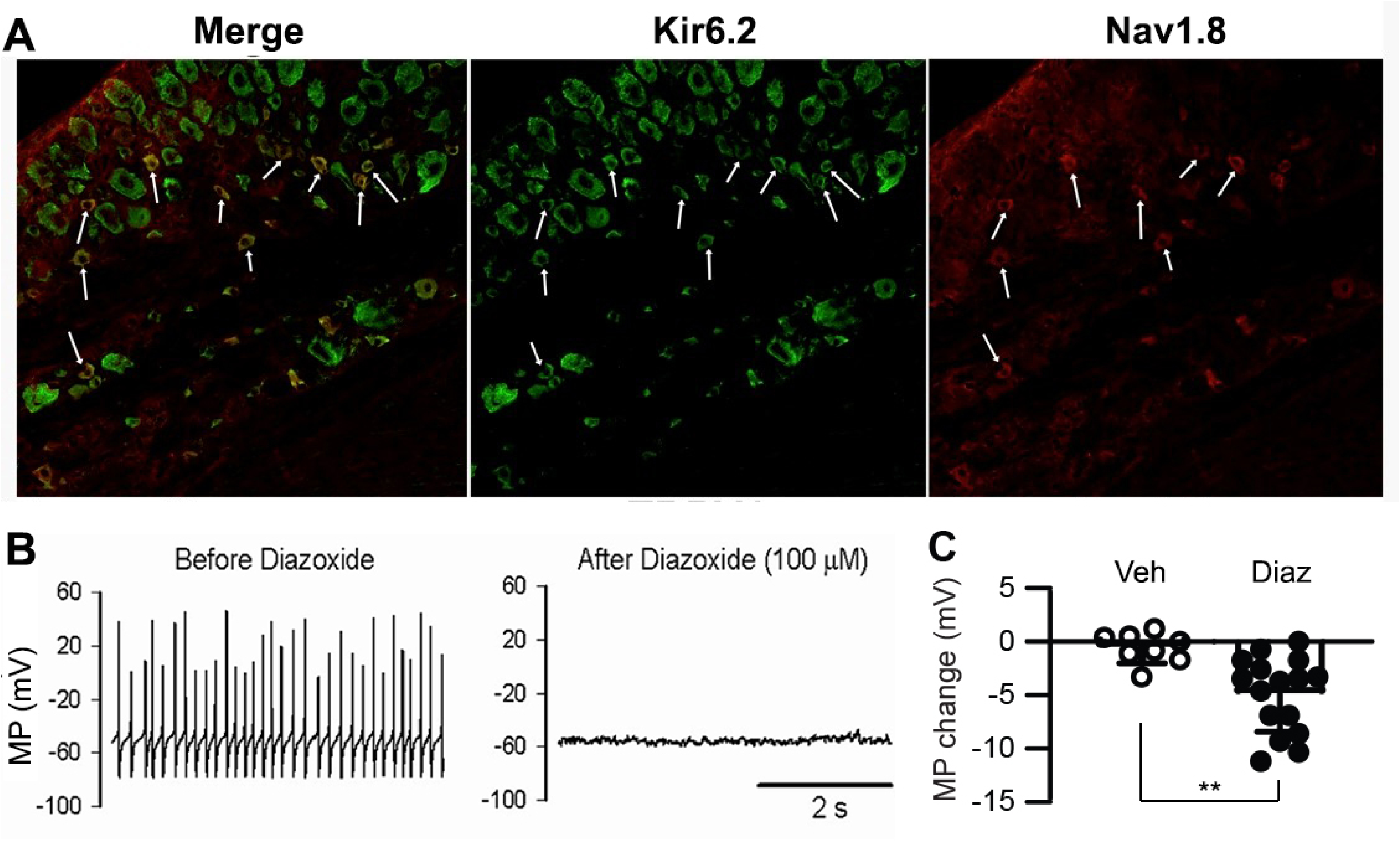
The effect of diazoxide on the excitability of DRG neurons. (**A**) Immunostaining of Kir6.2 in the DRG from naïve rat. (**B**) Representative recordings showing that the SA of DRG neurons from contusive animal were largely blocked after cells were exposed to diazoxide for 1 min. Note that cell membrane potential is hyperpolarized from about -51 mV to -55 mV. (**C**) Summary data indicates the changes in membrane potential upon local vehicle and diazoxide (100 μM) applications. **, P < 0.05, unpaired t test (two-tailed). Circle symbols in columns represent one DRG neuron each. Veh, vehicle. Diaz, diazoxide.

### 2. Diazoxide attenuates the development of chronic pain following SCI

Administering glibenclamide to block SUR1 immediately after spinal contusion reduces hemorrhagic necrosis, resulting in sustained improvements in locomotion and coordination ^3, 10^. However, this acute application exacerbates sensory dysfunction by affecting glial cell activation ^12^. Both neuronal hyperexcitation and glial activation contribute to secondary degeneration and the development of neuropathic pain after SCI ^21, 27–29^. Therefore, we investigate whether diazoxide improves sensory function after SCI with or without glibenclamide treatment.

At chronic stage of SCI (5 weeks post-contusion), we conducted mechanical and thermal sensitivity tests. In contrast to a prior study (Redondo-Castro, 2013), glibenclamide (administered 24 hours via an implanted Alzet pump, starting 30 minutes after impact at a rate of 200 ng/0.5 μl/h) alone did not exacerbate mechanical hypersensitivity tested by von Frey compared to the vehicle control group (**Fig. 2A**). However, it did worsen thermal (heat) hypersensitivity following SCI (**Fig 2B**). Repeated administration of diazoxide (from 3 days post-SCI for 10 days, 5 mg/kg, twice daily, i.p.) or vehicle was then applied. The hindpaw withdrawal threshold detected by von Frey test was reduced in both diazoxide alone group and the combination group (**Fig. 2A**). Additionally, diazxoide increased the latency of heat-induced hindpaw withdraw, and glibenclamide cannot reverse diazoxide-induced protective effect (**Fig. 2B**). In addition to evoked pain, SCI also induces spontaneous pain. We thus performed conditioned place preference (CPP) test to determine if diazxoide alters the development of SCI-induced spontaneous ongoing pain with or without glibenclamide. SCI animals treated with glibenclamide alone were more inclined to remain in the conditioned analgesic chamber (white chamber) than the vehicle control groups, suggesting that early glibenclamide application promotes the development of SCI-induced spontaneous pain. In contrast, diazoxide-treated rats, alone or in combination with glibenclamide, exhibited no preference for the white chamber, indicating that repeated diazoxide treatment during the acute phase mitigates later behavioral manifestations of spontaneous pain (**Fig. 2C**). While the combination treatment did not significantly increase the von Frey threshold compared to diazoxide alone, SCI animals received glibenclamide and diazoxide are more likely stay in the innately preferred black chamber. These findings suggest that early, repeated treatment of diazoxide with or without glibenclamide can prevent the development of SCI chronic pain.

**Figure 2.**
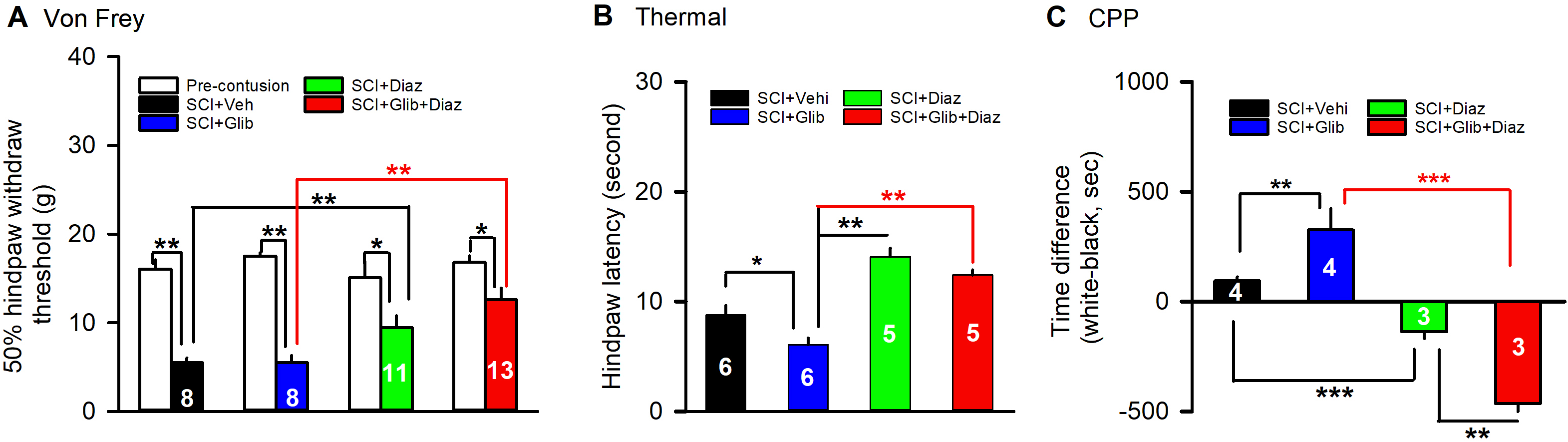
The effects of glibenclamide (starting 30 minutes after contusion for 24 hours) and diazoxide (starting 3 days post-SCI for 10 days) on development of pain behaviors after SCI. (**A**) Mechanical sensitivity of hindpaws was measured 5 weeks after initial traumatic spinal cord injury. Animal numbers were indicated on top of columns. * P < 0.05; ** P < 0.01. Two-tailed paired t-tests (before and after contusion of same group) or two-tailed unpaired t-tests (different treatment groups). (**B**) Thermal sensitivity of hindpaws was measured 5 weeks after spinal cord injury. Animal numbers were indicated in columns. * P < 0.05; ** P < 0.01. Two-tailed unpaired t-tests (different treatment groups). (**C**) Spontaneous pain was evaluated with CPP tests 42 days after contusion. Animal numbers were indicated in columns. ** P < 0.01; *** P < 0.001, One-way ANOVA followed by Turkey’s post hoc test. Veh, Vehicle; Glib, glibenclamide; Diaz, diazoxide.

### 3. Combining glibenclamide and diazoxide improves outcomes in motor recovery as compared to glibenclamide alone

Diazoxide and glibenclamide act as agonist and antagonist of SUR subunit, respectively. While glibenclamide improves the locomotor function following SCI, the impact of diazoxide on glibenclamide-induced beneficial effects is unknown. We thus tested the locomotor function of SCI rats after treatments.

Initially, SCI rats received either glibenclamide or its vehicle for 24 hours via an Alzet pump (200 ng/0.5 μl/h), starting 30 minutes post-impact ^3, 10^, Subsequently, diazoxide was administered for 10 consecutive days (i.p., 5 mg/kg, twice/day), commencing 3 days post-contusion. The effect of diazoxide on locomotor function post-SCI was assessed using the BBB (Basso, Beattie, and Bresnahan) 21-point open field locomotion scoring system. Consistent with previous reports ^3, 10^, glibenclamide significantly improved the BBB scores of the tested animals. However, diazoxide alone showed no improvement in BBB scores compared to the vehicle group from P7 to P35 (Fig 3A green dots). The combination of glibenclamide and diazoxide did not yield any additional improvement in BBB score compared to glibenclamide alone (Fig. 3A). It has been demonstrated that BBB scores exhibit a weak correlation with fiber conduction, particularly during the recovery stage when spontaneous remyelination is still enhancing ^24^. Hence, the horizontal ladder test was conducted to further assess the effect of treatments. Glibenclamide alone treatment significantly promoted horizontal ladder scores at tested time points post-SCI compared to vehicle control group. In contrast to the BBB scoring results, SCI rats treated with the combination of glibenclamide and diazoxide exhibited fewer hindlimb footslips compared to those receiving glibenclamide alone at day 35 post-SCI (**Fig. 3B**). These findings suggest that the combination of glibenclamide with diazoxide may enhance locomotor function after SCI.

**Figure 3.**
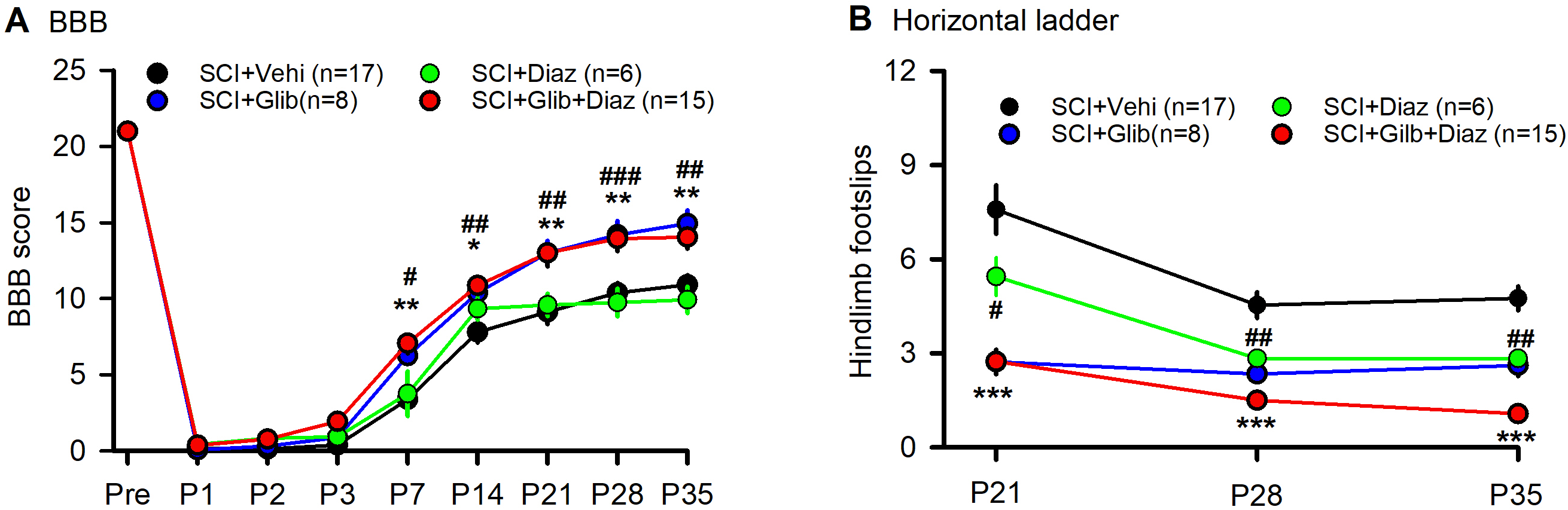
The effects of glibenclamide (starting 30 minutes after contusion for 24 hours) and diazoxide (starting 3 days post-SCI for 10 days) on development of locomotor behaviors after SCI. (**A**) BBB scoring was performed <u>-1</u>, 1, 2, 3, 7, 14, 21, 28, and 35 days after contusion. *, # P < 0.05; **, ## P < 0.01; ***, ### P < 0.001. *, **, **** Glibenclamide + Diazoxide vs Vehicle + Vehicle at same time points; #, ##, ### Glibenclamide + vehicle vs Vehicle + Vehicle at same time points. Two-tailed unpaired t-tests. (**B**) Horizontal ladder test was performed 21, 28, and 35 days after contusion. *** P < 0.001; ##, P < 0.05. **** Glibenclamide + Vehicle vs Vehicle + Vehicle at same time points; ##, Glibenclamide + vehicle vs Glibenclamide + Diazoxide at P35. Two-tailed unpaired t-tests.

Given that hemorrhage peaks between 3 to 24 hours post-acute SCI and resolves by day 7 ^18^, we test if delayed administration of diazoxide would yield better outcomes. Rats received glibenclamide (or its vehicle) for 24 hours via continuous intradermal infusion using an implanted Alzet pump (200 ng/0.5 μl/h), beginning 30 minutes post-impact ^3, 10^. Subsequently, diazoxide was administered for 7 consecutive days intraperitoneally (5 mg/kg, twice/day), commencing 6 days post-contusion. Similar as observed in early diazoxide groups (**Fig 2A** and **2B**), the delayed application of diazoxide significantly prevent the development of mechanical and thermal hypersensitivity even when combined with the acute glibenclamide (**Fig. 4A and B**). However, delayed diazoxide following glibenclamide did not further enhance the glibenclamide-induced locomotor improvement tested by BBB scoring and horizontal ladder tests (**Fig. 4C** and **4D** red bar). While delayed diazoxide alone reduced hindpaw foot slips, it did not improve BBB score after SCI (Fig 4C and 4D green bar).

**Figure 4.**
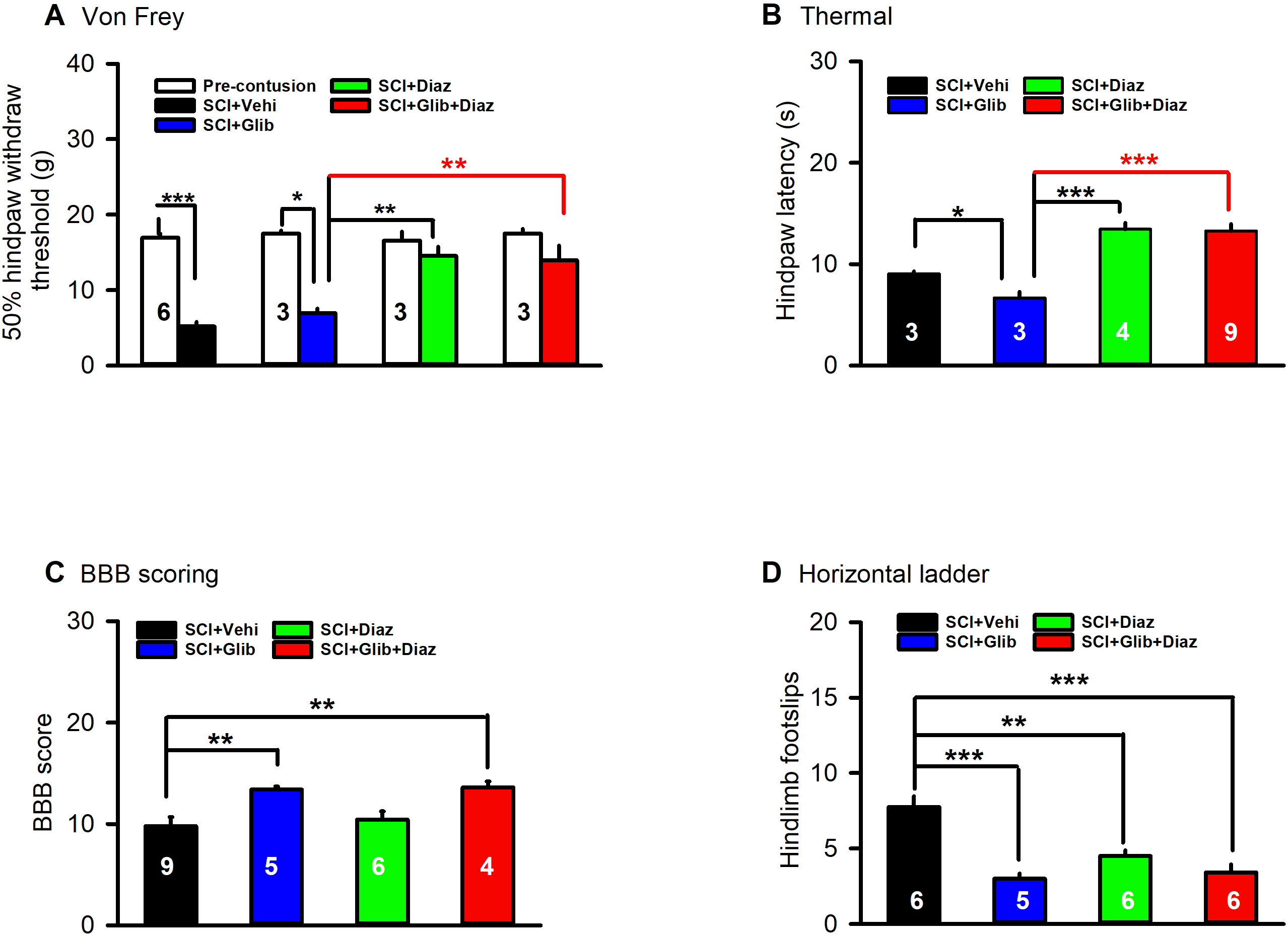
The effects of glibenclamide (starting 30 minutes after contusion for 24 hours) and diazoxide (starting 6 days post-SCI for 7 days) on development of locomotor and sensory behaviors after SCI. (**A**) Mechanical sensitivity of hindpaws was measured 35 days after initial traumatic spinal cord injury. Animal number for each group were indicated in columns. * P < 0.05; *** P < 0.001. Two-tailed paired t-tests (before and after treatment of the same group) and two-tailed unpaired t-tests (different groups). (**B**) Thermal sensitivity of hindpaws was measured 35 days after contusion. Animal number for each group were indicated in columns. * P < 0.05; *** P < 0.001. Two-tailed unpaired t-tests (different groups). (**C**) BBB scoring was performed 42 days after contusion. ** P < 0.01. One-way ANOVA followed by Turkey’s post hoc test. (**C**) Horizontal ladder test was performed 42 days after contusion. *** P < 0.001. One-way ANOVA followed by Turkey’s post-hoc test. Animal numbers were indicated in columns. Veh, vehicle; Glib, glibenclamide; Diaz, diazoxide.

### 4. Diazoxide alone or in combination with glibenclamide decreases the glial cell activation in the spinal cord after SCI

Following SCI, both astrocytes and microglia become activated in the spinal cord, contributing to locomotor and sensory dysfunction ^28, 30^. To assess the impact of treatments, we quantify the expression levels of GFAP and Iba-1 in the SCI spinal cord using Western blotting. We observed that glibenclamide alone significantly increased the expression levels of GFAP at the spinal cord compared to those in the vehicle-treated SCI groups. Conversely, diazoxide (starting 3 days after contusion for 10 days.) significantly decreased the expression level of spinal cord GFAP. Combining glibenclamide with diazoxide did not significantly further decrease GFAP compared to diazoxide alone (**Fig. 5A and B**). Both glibenclamide and diazoxide displayed no obvious effect on Iba-1 expression in the spinal cord following SCI.

**Figure. 5.**
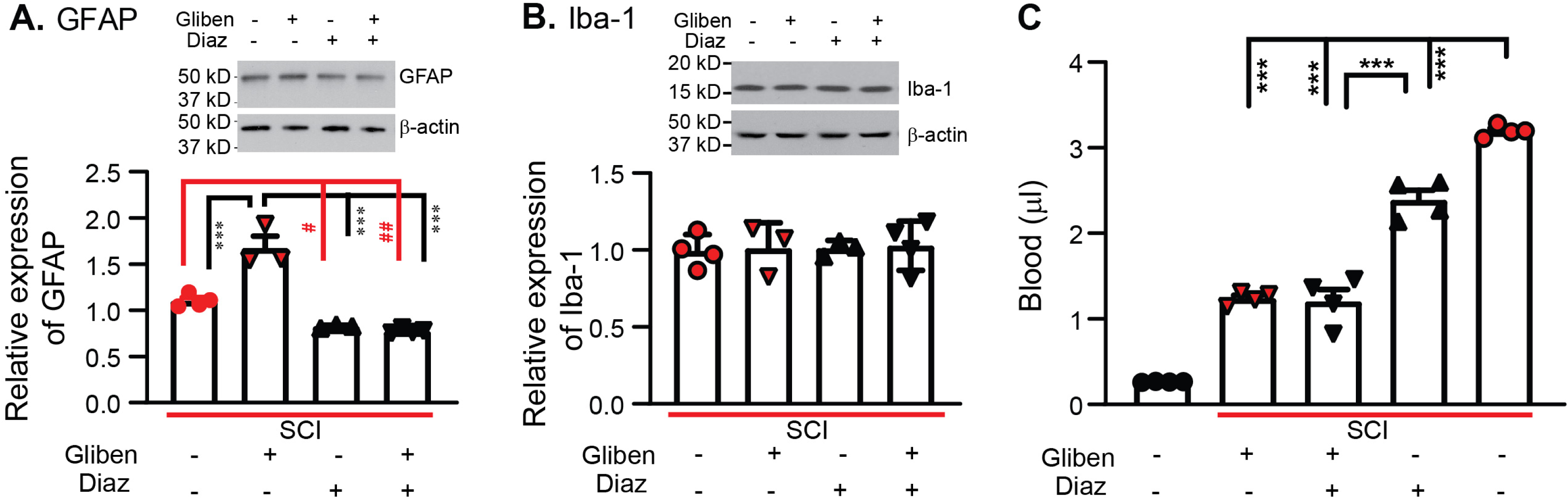
The effects of glibenclamide (starting 30 minutes after contusion for 24 hours) and diazoxide (starting 3 days post-SCI for 10 days) on SCI-induced activation of glial cells and hemorrhage in the spinal cord. (**A**) Quantification of GFAP expression normalized to β-actin in L4/L5 spinal cords. # P < 0.05, ## P < 0.01, *** P < 0.001. One-way ANOVA followed by Turkey’s post-hoc test. (**B**) Quantification of Iba-1 expression normalized to β-actin in L4/L5 spinal cords. (D) Quantification of extravascular blood in the spinal cord lesion site. *** P < 0.001, One-way ANOVA followed by Turkey’s post-hoc test. Filled symbols in each column represents individual animals. Gliben, glibenclamide; Diaz, diazoxide.

### 4. Diazoxide decreases the extravascular blood after SCI

While the SUR1 antagonist glibenclamide has been shown to reduce hemorrhage following SCI ^3, 6^, we tested whether the SUR1 agonist diazoxide exacerbates hemorrhage after SCI. SCI rats received glibenclamide (or its vehicle) for 24 hours via an implanted Alzet pump, starting 30 minutes after contusion, followed by diazoxide for 1 day intraperitoneally (5 mg/kg, twice daily), starting 3 days post-contusion. Spinal cords encompassing the contusion site were collected on day 4. In contrast to the spinal cords from naïve rats, extravascular blood detected in the spinal cords subjected to contusion was significantly increased. Same as previous reports ^3^, the spinal cord hemorrhage was significantly reduced by acute glibenclamide infusion, as well as by diazoxide alone during the subacute stage, although the effect induced by diazoxide was relatively weaker than that induced by glibenclamide. Nevertheless, the combined application of SUR1 antagonist and agonist at different stage did not further reduce spinal cord hemorrhage compared to glibenclamide alone (**Fig. 5C**).

### 5. Diazoxide does not further preserve spinal cord tissue after glibenclamide treatment

Histological evidence was gathered to assess the effect of different treatments on spared tissue, particularly by measuring the volume of spared tissue in the spinal cord (gray matter and white matter) using eriochrome cyanine (EC) staining. There was greater surviving spinal tissue in the glibenclamide-treated SCI animals compared to that in the vehicle-treated rats at P42 (Fig 6, black and blue bar). Diazxoide alone also increased the spared white matter, but not gray matter, compared to the SCI vehicle control group (Fig. 6 green and black bar). However, no significant differences in spared tissue of both gray and white matter were observed between the group with combination of drugs and the group with glibenclamide alone (**Fig. 6**, red and blue bar). These findings suggest that glibenclamide, rather than diazoxide, plays a critical role in decreasing the degeneration of spinal tissues, especial gray matter, following SCI.

**Figure 6.**
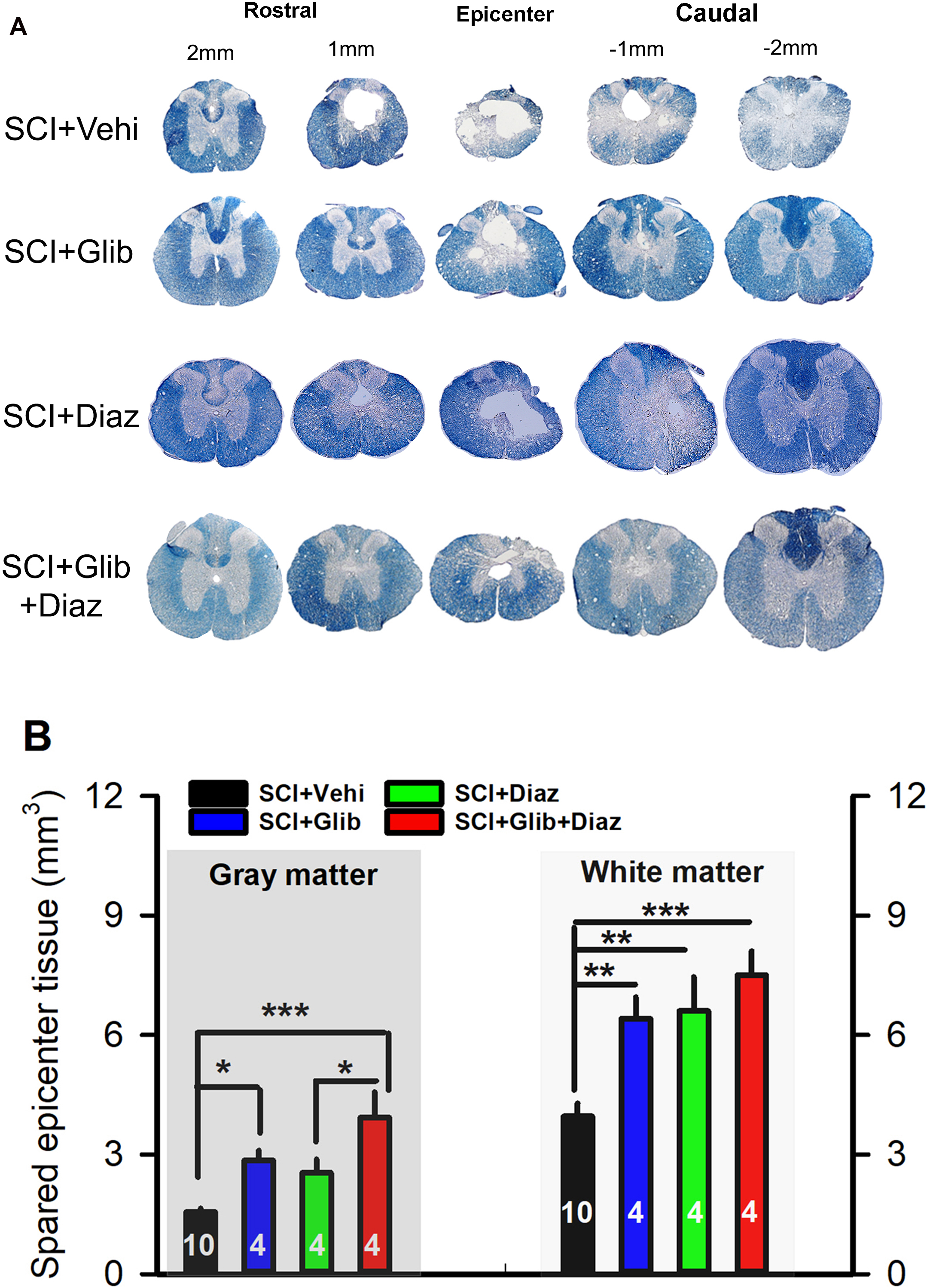
The effects of glibenclamide (starting 30 minutes after contusion for 24 hours) and diazoxide (starting 3 days post-SCI for 12 days) on SCI-induced tissue loss in the spinal cord. (**A**) Representative images showing EC staining of spinal cord sections of treated SCI rats. (**B**) The graph indicates the quantification of the spared gray matter and white matter of spinal cords at the epicenter of contusion site from <u>glibenclamide and/or diazoxide-treated SCI rats</u>. * P < 0.05, one-way ANOVA followed by Turkey’s post hoc test. Veh, vehicle; Glib, glibenclamide; Diaz, diazoxide.

## Discussion

Spinal cord injury results in diverse impairments, encompassing motor dysfunction and chronic pain. Both neuronal hyperactivity and glial activation are implicated in secondary degeneration post-SCI and the onset of neuropathic pain ^21, 27–29^. Although blocking SUR1 with glibenclamide immediately after spinal contusion yields sustained improvements in locomotion and coordination ^3, 10^, it exacerbates sensory dysfunction, potentially by modulating glial cell activation ^12^. K_ATP_ channels, comprising Kir6.1 or Kir6.2 subunits and SUR, are widely distributed in both glial cells and neurons ^7–9^, exerting regulatory control over glial activation and neuronal excitability. Our findings suggest that timely modulation of SUR subunit activity, through its agonist and antagonist, enhances both motor and sensory function following SCI.

A previous study indicated that administering glibenclamide alone during the acute stage exacerbates the progression of mechanical pain, as assessed by the Randall Selitto test following SCI ^12^. Although our thermal and conditioned place preference tests showed a similar trend, von Frey tests indicated that glibenclamide did not further decrease the hindpaw withdrawal threshold. The discrepancy between von Frey and Randall Selitto tests lies in their methodologies: the von Frey test evaluates cutaneous allodynia, likely mediated by Aβ-fibers^31^, whereas the Randall Selitto test likely gauges non-cutaneous hypersensitivity transmitted through nociceptors. Thermal hypersensitivity and spontaneous pain are mediated by nociceptors ^21, 32^. While both cutaneous allodynia and non-cutaneous hypersensitivity arise after SCI ^21, 32^, glibenclamide might primarily augment the plasticity of nociceptors.

After spinal cord injury, both astrocytes and microglial cells become activated not only at the lesion site but also in areas both above and below the level of the injury, contributing to the development of SCI-induced neurobehavioral dysfunction^28, 30^. K_ATP_ channels are expressed in both astrocytes and microglia ^9, 33^. While the expression levels of astrocyte marker GFAP in the lesion site is significantly modulated by glibenclamide and diazoxide, changes in the expression levels of the microglial marker Iba-1 are not significant. Microglia become activated in the spinal cord on day 3 post-SCI^34^. Iba-1-positive microglia can be classified into two opposing subtypes: classical proinflammatory (M1) and polarized anti-inflammatory alternatives (M2), which have both harmful and beneficial effects in neurodegenerative diseases^35, 36^. Microglia in the spinal cord play a crucial role in both the initiation and maintenance of SCI-induced chronic pain ^28, 29^. While we observed a protective effect of diazoxide in the development of chronic pain, it is possible that diazoxide enhances the transition of microglia from M1 to M2 without changing in Iba-1 expression level^33^. This possibility requires further evaluation.

Both glibenclamide and diazoxide decrease extravascular blood in the spinal cord. Hemorrhage peaks at 3 to 24 hrs after acute SCI, and dissolved by day 7 ^18^. Acute administration of glibenclamide reduces secondary hemorrhage in the spinal cord after SCI by preventing fragmentation of spinal cord capillaries through TRPM4 ion channels ^5^. The absorption of hematomas after hemorrhage primarily relies on the phagocytes’ ability to engulf red blood cells ^37^. Activation of both K_ATP_ channels and TRPM4 channels contributes to enhanced activity of monocytes/macrophages ^38, 39^, it is thus possible that diazoxide mitigates extravascular blood by promoting hematoma absorption. However, the impact of delayed administration of diazoxide on this effect may not be significant in combination treatment, as the pool of extravascular blood is already limited by acute glibenclamide application.

Heightened neuronal excitability and activation of spinal cord glial cells are pivotal in the onset of dysfunction following SCI ^28, 29, 40^. Targeting neuronal hyperexcitability with retigabine, a selective KCNQ ion channel activator, has shown improvements in both motor and sensory dysfunction after SCI ^27^. Intraperitoneal injection of minocycline, a nonspecific microglial inhibitor ^41^, reduces lesion size and hyperexcitability of spinal neurons in a rodent SCI model. Diazoxide enhances the activity of K_ATP_ channels expressed in both neurons and glial cells, mitigating both neuronal excitability and glial activation, thereby improving sensory function with or without concurrent glibenclamide administration. When comparing the early application of diazoxide after glibenclamide with delayed application, similar outcomes are observed for sensory function but not for motor function. Early diazoxide administration, but not delayed administration, further enhances glibenclamide-promoted locomotor function, as evidenced by horizontal ladder tests. This study suggests that neuronal hyperexcitation and/or spinal inflammatory activity during the acute stage are more crucial for the development of motor dysfunction than the pain chronification induced by SCI.

## Data Availability Statement

The data that support the findings of this study are available from the leading corresponding author upon reasonable request.

## Author Contributions

Participated in research design: Q. Yang and Z. Wu

Conducted experiments: Z. Wu, Q. Yang, G. Xu, C. Prater, M. Maskey

Performed data analysis: Z. Wu, Q. Yang, G. Xu

Wrote or contributed to the writing of the manuscript: Q. Yang and Z. Wu

## Conflicts of Interest

The authors declare no conflict of interest.

## Footnotes

The research was funded by grants from National Institutes of Health (NIH) grants (CA208765 to Q.Y., DC016328 to Z.W.), the Craig H. Neilsen Foundation (383428 and 993497 to Q.Y.), and CDMRP SCIRP (W81XWH-14-1-0593 to Q.Y.).

## List of Nonstandard Abbreviations

BBB: Baso, Beattie, Bresnahan
CPP: Conditional place preference
EC: Eriochrome Cyanine
K_ATP_: ATP-gated potassium channels
PHN: Progress hemorrhagic necrosis
SCI: Spinal cord injury
SD: Sprague-Dawley
SUR: Regulatory sulfonylurea
TRPM4: Transient receptor potential cation channel subfamily melastatin-4

## Notes

**Conflict of interest statement:** The authors have declared that no conflict of interest exists.

### Competing Interest Statement

The authors have declared no competing interest.

